# A Metric for Quantifying the Resolution of Molecular Assays

**DOI:** 10.1101/2021.05.07.443139

**Authors:** Brandon D. Wilson, Michael Eisenstein, H. Tom Soh

## Abstract

Many assay developers focus on limit of detection (LOD) as a primary performance metric, and LOD is indeed useful for assays designed to determine the presence or absence of an analyte. However, LOD is less useful for ‘continuous assays’ designed to discriminate between concentrations of an analyte—*e*.*g*., glucose monitoring in diabetes, where clinical care is guided by particular concentration ranges and thresholds. In such scenarios, it would be valuable to quantify discriminatory resolution—*i*.*e*., whether an assay can differentiate 1 pM from 10 pM— but no such standardized metric currently exists. Here, we propose a useful solution, termed ‘resolution of molecular concentration’ (RMC). RMC offers a simple means for characterizing quantitative resolution and quickly comparing the quantitative performance of assays. By raising awareness of the limitations of current metrics for evaluating assay performance, we hope to empower the molecular diagnostics community to evaluate their methods in a more application-appropriate manner.

## Main

Assays for measuring molecular concentrations are indispensable in diagnostics, drug development, and many other areas of basic and clinical research. Although there are many metrics for evaluating the analytical performance of an assay, perhaps none is used more widely than the limit of detection (LOD)—the lowest concentration that can be statistically differentiated from background signal (**Box 1**).^1^ LOD is commonly used to distill the performance of assays into a single metric that enables head-to-head comparisons.

Although a low LOD is often treated as the most important aspect of an assay’s performance and a primary goal for assay developers, it is not the most useful metric for all molecular detection applications. On one hand, LOD is very useful for describing the clinical utility of assays whose goal is to determine the presence or absence of a certain molecule (*e.g*. infectious disease testing). On the other hand, LOD is less useful for ‘continuous assays’ that are intended to distinguish between different concentrations of an analyte. An assay with an LOD of 23 nM for glucose marks an impressive technical feat^2^, but this metric is immaterial to whether the assay can differentiate a healthy ∼8 mM glucose concentration from the ∼12 mM glucose concentration found in a diabetic patient. In such cases, the ability to accurately resolve changes in concentration is far more relevant than the ability to distinguish a given concentration from background signal.

Unfortunately, there is currently no standardized way to describe an assay’s discriminatory power in terms of distinguishing one concentration from another. A variety of metrics have been proposed for describing the quantitative performance of molecular assays^3^, and many authors have discussed mechanisms for assessing the accuracy or precision of a single measurement^4,5^. One set of parameters that is used occasionally is the lower limit of quantification (LLOQ) and the upper limit of quantification (ULOQ). These metrics have been described as the range of target concentrations over which precision is greater than a certain threshold, or alternatively, may be calculated in the same manner as LOD but with a cutoff of 10 standard deviations rather 3. Regardless of the definition, LLOQ and ULOQ deal with the precision of measurements at a single concentration, rather than facilitating comparison *between* concentrations. Similarly, precision, which is usually expressed as the coefficient of variation (CV) at a given concentration, refers only to the repeatability of a single measurement. Determining that an assay has a CV of 10% at 1 µM target concentration does not indicate whether the assay can reliably distinguish between 1 µM and 2 µM of that target.

In this work, we propose a new analytical framework called resolution of molecular concentration (RMC), which offers a straightforward and physiologically relevant means to characterize an assay’s quantitative resolution and quickly compare quantitative performance across assays. Our goal here is to incite discussion and raise awareness of the limitations to the metrics we use to evaluate and select assays for their intended applications. Lastly, by providing resources for researchers to easily implement RMC, we hope to guide better assay selections and ultimately increase their clinical utility.

### Box 1

**Brief Review of LOD**

Molecular quantification ultimately involves three basic steps: 1) a defined chemical system that converts an unknown concentration into an observable signal; 2) a standard curve derived from known concentrations of target that defines the functional relationship between concentration and signal; and 3) an inverse function derived from the standard curve that converts signal back to concentration. The performance of this process is often evaluated in terms of the limit of detection (LOD): the lowest concentration that can be statistically distinguished from background signal. LOD is typically calculated as the concentration that gives a signal equal to the background signal plus three standard deviations:

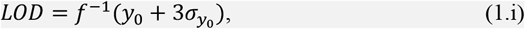

where *f*^−1^ is the inverse of the standard curve function. In the linear example below (**Figure i**), a standard curve defines the functional relationship between concentration and signal. *f*^−1^ evaluated at the background signal plus three standard deviations defines the LOD.

**Figure i.**
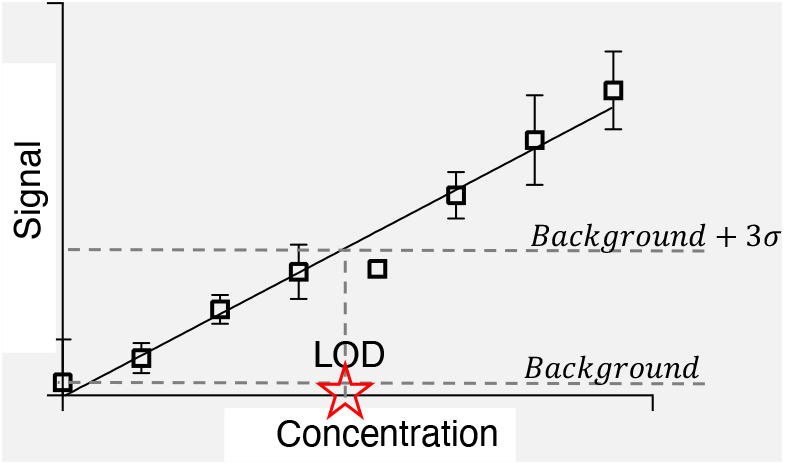
Determining LOD from a standard curve. LOD is conventionally calculated as the concentration that yields a signal equivalent to the background signal plus three standard deviations, as predicted by the standard curve.

LOD is straightforward to calculate in this linear example:

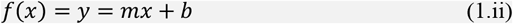

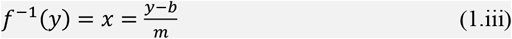

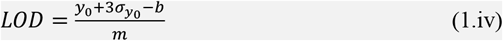

## Resolution of molecular concentration (RMC)

RMC is designed to identify the smallest fold-change in target concentration that can be distinguished with > 99.5% certainty. To illustrate this, imagine an arbitrary assay where we want to compare two concentrations, *x* and *μx*, which differ by *μ*—a scalar constant of proportionality greater than 1 (**Figure 1**). The smallest value of *µ* for which the resultant signals *S*_1_ =*f*(*x*) and *S*_2_ =*f*(*μx*) are statistically distinguishable is the basis for RMC.

**Figure 1.**
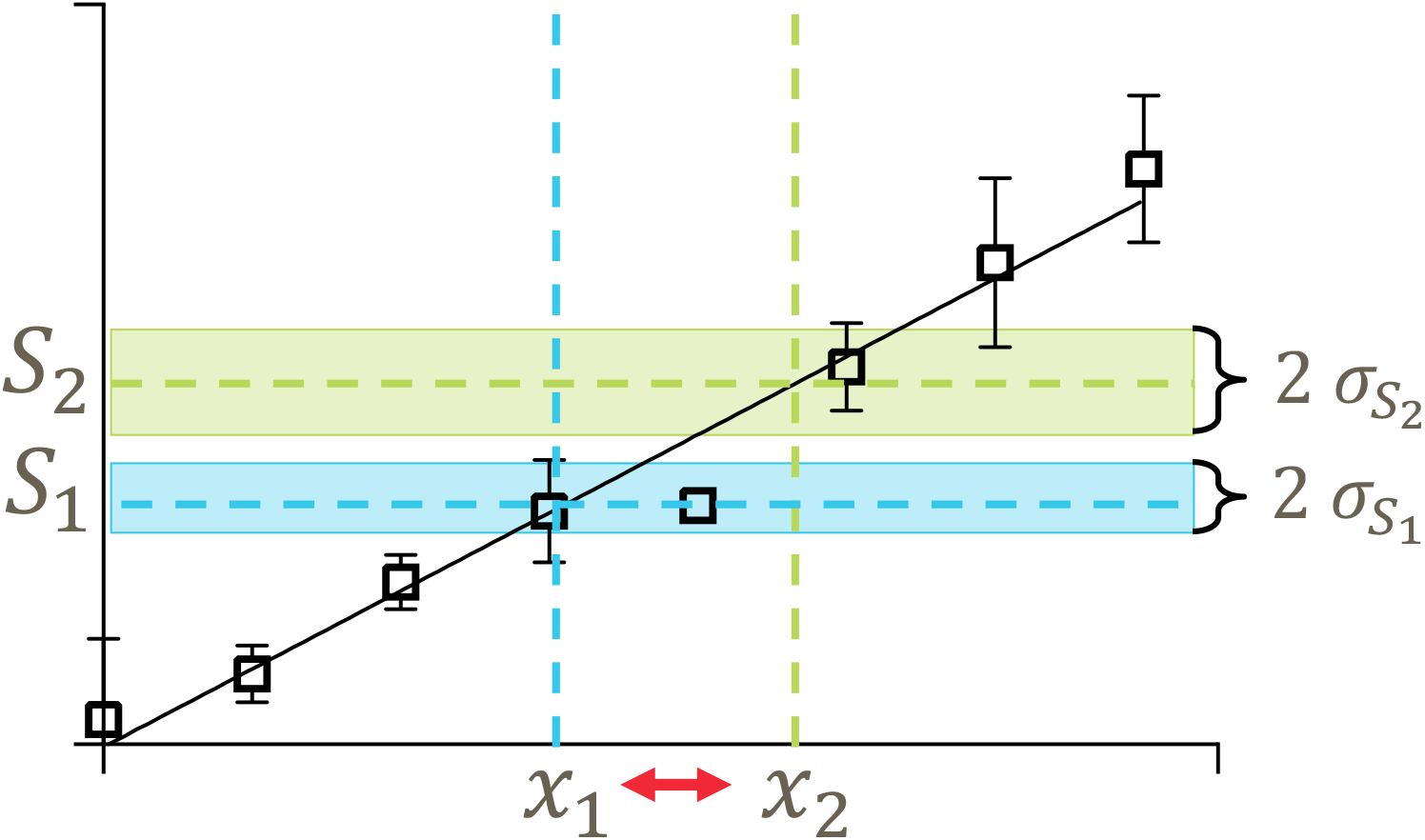
Schematic of resolution. If we consider an arbitrary assay where there is a concentration-dependent signal, how do we know whether the resultant signals from two different concentrations are meaningfully distinguishable? We can use a t-test to determine whether these two signals are distinguishable with 95% certainty. To translate this into a tangible metric of resolution, we let *x*_1_ =*x* and *x*_2_ =μ*x*, and solve for µ as a function of *x* (eq. 4). Now µ(*x*) represents resolution in terms of the smallest fold change in concentration that can be distinguished with 95% certainty.

The certainty with which we can distinguish two concentrations is ultimately a function of how confident we are in the parameterization of the standard curve. To quantify this, we perform a t-test between resultant signals and solve for the smallest *μ* that meets a defined threshold of statistical certainty. Equation 1 is based on an arbitrary threshold of 99.5% certainty—equivalent to approximately three standard deviations, in keeping with the colloquial LOD definition—and the assumption that the hypothetical measurements S_1_ and S_2_ are derived from triplicate measurements.

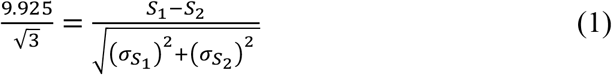

The expected signal can be estimated from the standard curve, which is a function of target concentration *x* (**eq. 2**), and a known set of fit parameters {*c*_1_, *c*_2_, *c*_3_, ⃛} that define the standard curve function (*e.g*., *B*_*max*_, *K*_*D*_, *y*_0_) for a standard Langmuir fit (**eq. 3**).

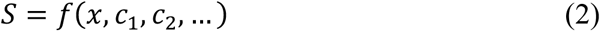

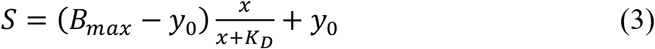

In assessing the relationship between two concentrations of interest, *x* and *μx*, equation 1 becomes:

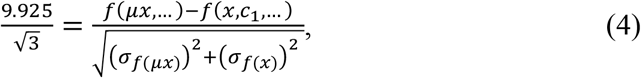

where *σ*_*f*(μ*x*)_ and *σ*_*f*(*x*)_ are calculated via propagation of errors:

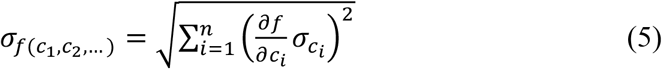

The RMC function is determined by solving **eq. 4** for *μ* as a function of *x. μ*(*x*) represents the smallest fold-change that can be discriminated from concentration *x* with 99.5% certainty. Similarly, *μ*(*x*) * *x* is the lowest concentration that is differentiable from *x* with 99.5% certainty. For instance, *μ*(2 *mM*) =1.5 would mean that an assay can discriminate between the signals generated by 2 mM and 3 mM target with 99.5% certainty. We note that *μ*(*x*) is a function of target concentration, and the shape of *μ*(*x*) is highly predicated on the shape of the binding curve. For instance, in a standard Langmuir binding curve (**eq. 3**), we expect *μ* to adopt a minimum near *x* =*K*_*D*_. In the following sections, we illustrate the utility of *μ*(*x*) through two case studies.

### Case study #1 | Choosing the right ELISA

Enzyme-linked immunosorbent assays (ELISAs) are the most commonly used assay format for the detection of proteins. In many cases, there are multiple ELISA kits available for the same analyte. It is therefore important to be able to compare performance to be able to pick the best assay for the given clinical context. While this comparison is traditionally made by comparing LODs, we demonstrate here how one might instead use RMC to pick the best assay for quantifying C-reactive protein (CRP), a biomarker of inflammation that has been demonstrated to be predictive of certain categories of cardiovascular disease^6^.

Stratification of patient risk for adverse cardiac events requires distinguishing relatively small (∼2-fold) changes in CRP concentration^6^, where 0.3 µg/mL, 0.6 µg/mL, 1.5 µg/mL, 3.5 µg/mL, and 6.6 µg/mL represent the 10^th^, 25^th^, 50^th^, 75^th^ and 90^th^ percentile, respectively. Since a two-fold change in concentration correlates to a ∼25±12% increase in risk of cardiovascular disease^6^, clinicians would be particularly interested in an assay that could stratify patients into these risk categories based on CRP concentration. We use RMC to evaluate three commercial ELISAs (**Figure 2a**) for their suitability for this application, with LODs of 0.022, 0.003, and <0.01 ng/mL for assay A, assay B, and assay C, respectively (**Figure 2b**).

**Figure 2.**
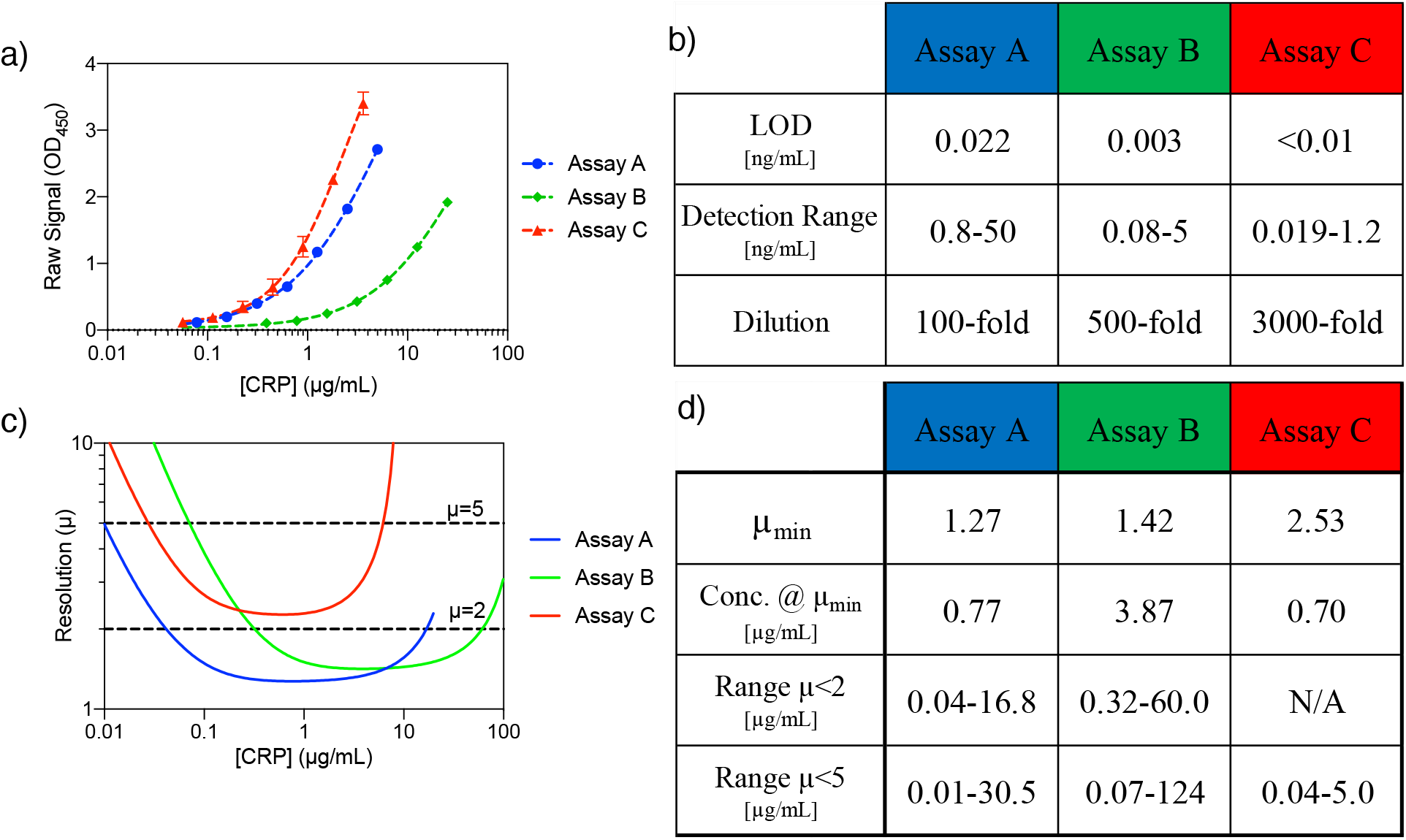
Comparing commercial assays for CRP. **(a)** The standard curves reported in the manufacturer datasheets were converted to CRP concentrations from undiluted samples by multiplying by the relevant dilution factors. Error bars represent standard deviations for two replicates. **(b)** Analytical performance values of the three commercial ELISA kits as reported by the manufacturers. **(c)** Plot of resolution in terms of μ vs concentration for each assay. **(d)** Analytical parameters of the assays that can be obtained from the RMC analysis. Assay information: Assay A is manufactured by R&D Systems (Cat. No. DCRP00), Assay B is manufactured by Invitrogen (Cat. No. BMS288INST), and Assay C is manufactured by Invitrogen (Cat. No. KHA0031).

Based on the clinical parameters described above, our ideal assay would have *μ* < 2 over the range of concentrations from 0.30 to 6.6 µg/mL. To calculate RMC, we performed four-parameter fits to these standard curves and solved for *µ(x)* (**Figure 2c**). The RMC function produces rich information about each assay. For instance, we can compare the peak resolutions of the assays. Assay A achieves a maximum resolution of *µ* = 1.27 when [CRP] = 0.77 µg/mL; this means that the smallest difference in concentration that can be resolved with 99.5% confidence is a 27% change from 0.77 to 0.98 µg/mL. Assay C achieves peak resolution for a 153% change in concentration from 0.70 to 1.8 µg/mL, whereas assay B can resolve a 42% change in concentration from 3.9 to 5.5 µg/mL. None of this information could be gleaned from the LOD alone.

Importantly, we can use this novel information to make better assessments of how an assay will work in a particular application. Although assay C has a very low reported LOD of < 0.01 ng/mL, it also yields greater uncertainty in terms of the calculated CRP concentration, and therefore has the lowest quantitative resolution. Assay A, on the other hand, is capable of reliably stratifying patients into different risk categories with high statistical certainty over the entire range of clinical concentrations. This assay also outperforms assay B in terms of resolution, even though the latter has the lowest reported LOD of the three tests.

This analysis can similarly be used to evaluate assay performance for the detection of sepsis. Baseline CRP concentration is below 10 µg/mL in 99% of individuals,^7^ and sepsis typically results in CRP concentrations of over 50 µg/mL^8^. Therefore, the ideal assay will have *μ* < 5 over the concentration range of 10–50 µg/mL. The resolution curves (**Figure 2c**) suggest that assays A and B would both be up to the task. However, since many immunoassays suffer from the hook effect, where high concentrations of target actually result in a *decrease* in signal,^9^ and since assay A has only been validated at up to ∼5 µg/mL CRP, it would be necessary to confirm that assay A does not experience the hook effect at higher concentrations. Therefore, assay B would be the best choice for monitoring the development of sepsis, offering a *μ* < 2 throughout the relevant concentration range.

### Case study #2 | Comparing technologies for therapeutic drug monitoring

Individual differences in metabolism and pharmacokinetics can result in drug underdosing or overdosing, which are two of the primary modes of failure in clinical trials^10^. RMC could also play a key role in this context for analyzing technologies that support therapeutic drug monitoring (TDM), which would enable researchers and physicians to provide personalized doses that stably maintain desired drug concentrations within even the narrowest therapeutic window. However, it is essential that TDM assays or devices are capable of statistically resolving concentrations at the borders of the toxic-to-therapeutic range, and LOD and other metrics do not currently address this important factor.

RMC is particularly well suited to tackle this problem, which we demonstrate by looking at four commercial point-of-care assays that Genentech researchers have previously compared for their ability to quantify circulating concentrations of monoclonal antibody therapeutic-A (Anti-A)^11^. The four assays include two signal-on (assays D and E) and two signal-off assays (assays F and G), feature a variety of readouts (*e.g*., electrochemical (assays D and E), piezoelectric (assay F), and enzymatic (assay G)), and exhibit detection ranges spanning three orders of magnitude (**Figure 3a**).

**Figure 3.**
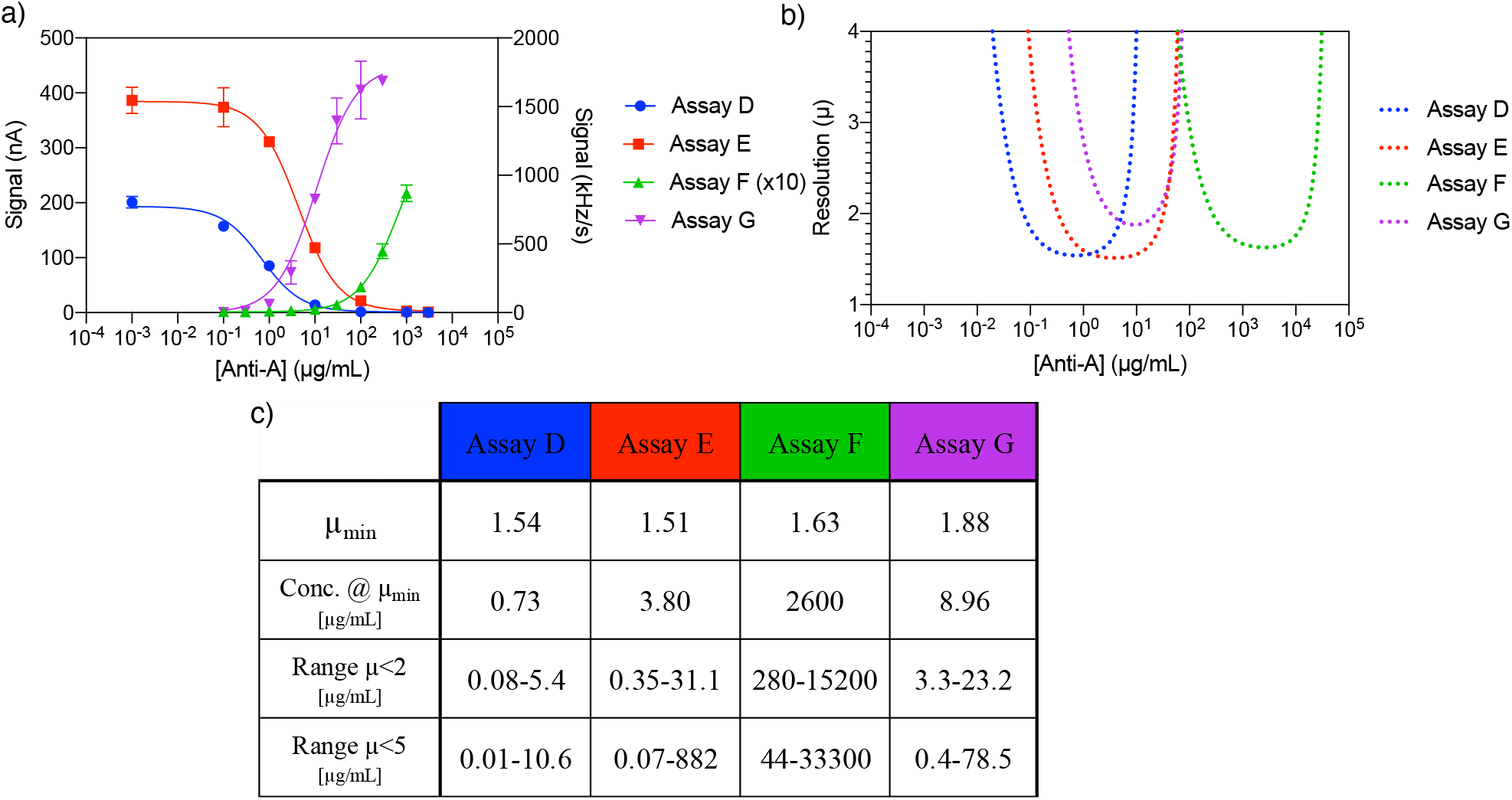
Point-of-care assays for therapeutic drug monitoring. **(a)** We looked at four assays for monitoring circulating concentrations of Anti-A^11^. Assays F and G are signal-off (plotted on the left-hand y-axis), while assays D and E are signal-on (plotted on the right-hand y-axis). The readout values of assay F have been multiplied by 10 to fit on the same axis as assay G. **(b)** Resolution values for the four point-of-care assays. **(c)** Analytical parameters of the assays related to resolution. Following the nomenclature of reference ^11^: Assay D is Proxim S1, Assay E is Proxim S2/3, Assay F is Qorvo Direct, and Assay G is Qorvo Enzyme.

RMC analysis (**Figure 3b**) quickly reveals that these four assays have very different performances and would therefore be most useful in different clinical scenarios. For instance, the concentrations that produce the greatest quantitative resolution are 0.73 µg/mL for assay D, µ_min_=1.54, versus 2,600 µg/mL for assay F, µ_min_=1.63 (**Figure 3c**). We are also able to define ranges of concentrations over which each assay achieved a certain threshold of quantitative resolution—we used µ < 2 as an example in this analysis and observed that the lowest concentration range for which µ < 2 was 0.08–5.36 µg/mL for assay D, whereas the highest concentration range was 280–15,200 µg/mL for assay F (**Figure 3c**). Notably, there is a gap in the dynamic ranges of these four assays: none of the assays can reliably resolve concentrations between 21 and 280 µg/mL. Thus, RMC analysis would enable clinicians and researchers to choose assays and devices that are best suited to the clinical scenario and desired drug dosage range at hand.

## Conclusion

Contemporary research continues to rely heavily on LOD as the primary descriptor of assay performance. Even though LOD provides important information about how well an assay can distinguish target concentration from background, it offers little insight into how well an assay can differentiate one target concentration from another. Here, we have explored the statistical certainty with which one concentration can be differentiated from another, with RMC—a metric that is easy to understand and appropriately characterizes the quantitative resolution of an assay. RMC is described in terms of *μ*(*x*), a function that represents the smallest fold-change that can be discriminated from concentration *x* with 99.5% certainty.

RMC reveals important information about assay utility that cannot be derived from LOD alone and enables researchers to quickly compare the quantitative power of two assays. RMC can be applied to virtually any continuous assay independent of the underlying detection mechanism and used to guide the selection of assays for diagnostics, therapeutic dosing, and other scenarios in which discriminatory power is critical. It is our hope that the concepts presented herein will empower the molecular diagnostics community by adding a more relevant dimension for evaluating assays in an application-appropriate manner.

## Acknowledgements

We thank Marius Tirlea and Michael Sklar of the Stanford Statistics Group for their helpful discussion on the manuscript.

